# Can DyeCycling break the photobleaching limit in single-molecule FRET?

**DOI:** 10.1101/2022.02.08.479542

**Authors:** Benjamin Vermeer, Sonja Schmid

## Abstract

Biomolecular systems, such as proteins, crucially rely on dynamic processes at the nanoscale. Detecting biomolecular nanodynamics is therefore key to obtaining a mechanistic understanding of the energies and molecular driving forces that control biomolecular systems. Single-molecule fluorescence resonance energy transfer (smFRET) is a powerful technique to observe in real-time how a single biomolecule proceeds through its functional cycle involving a sequence of distinct structural states. Currently, this technique is fundamentally limited by irreversible photobleaching, causing the untimely end of the experiment and thus, a prohibitively narrow temporal bandwidth of ≤ 3 orders of magnitude. Here, we introduce ‘*DyeCycling*’, a measurement scheme with which we aim to break the photobleaching limit in single-molecule FRET. We introduce the concept of spontaneous dye replacement by simulations, and as an experimental proof-of-concept, we demonstrate the intermittent observation of a single biomolecule for one hour with a time resolution of milliseconds. Theoretically, DyeCycling can provide >100-fold more information per single molecule than conventional smFRET. We discuss the experimental implementation of DyeCycling, its current and fundamental limitations, and specific biological use cases. Given its general simplicity and versatility, DyeCycling has the potential to revolutionize the field of time-resolved smFRET, where it may serve to unravel a wealth of biomolecular dynamics by bridging from milliseconds to the hour range.

## 1 Introduction

Biomolecular nano-dynamics are essential for all life on earth. In Richard Feynman’s words: “Everything that living things do can be understood in terms of the jiggling and wiggling of atoms” [1]. Nevertheless, our understanding of how biological function arises at the nanoscale has remained surprisingly modest. In fact, it is still challenging to experimentally observe the specific structural transitions that single biomolecules undergo during their functional cycle. Single-molecule Fluorescence (or Förster) Resonance Energy Transfer (smFRET) is one of the most popular techniques capable of observing the time evolution of one single biomolecule through various functional states [2–4]. However, until now, smFRET has been fundamentally limited by irreversible photo-bleaching, causing poor temporal bandwidths as reviewed below.

Here, we present DyeCycling, a measurement scheme designed to break the photo-bleaching limit in smFRET, and thus to observe a single molecule for up to hours with a time resolution of milliseconds. We first motivate DyeCycling by discussing the broad range of biomolecular dynamics using the example of proteins, and introduce smFRET to the non-expert. We summarize the current state of surface-immobilized smFRET, and review the achievable temporal bandwidths. Next, we present the concept of DyeCycling using simulations, and discuss various experimental implementations. We provide an experimental proof-of-concept of DyeCycling, and discuss its benefits and current experimental as well as ultimate fundamental limits. Lastly, we highlight specific biological use cases that will benefit from the increased observation time accessible via DyeCycling.

### 1.1 The importance of protein dynamics and the energies involved

The specific biological function of a protein is encoded in its amino-acid sequence and post-translational modifications, which define its molecular shape and its flexibility in aqueous solution [5]. Beyond intra-molecular dynamics, proteins undergo (often transient) inter-molecular interactions, adding a second dynamic component to protein function. Thanks to great progress in structural biology, static protein 3D structures with atomic resolution have become readily available mainly by x-ray crystal diffraction and electron microscopy. After 50 years of collaborative efforts, the Protein Data Bank (PDB) now holds over 185’000 macromolecular structures [6] – a great resource of information that is now harnessed by AlphaFold2, an artificial-intelligence approach to protein structure prediction [7,8]. However, comparable protein dynamics resources remain missing, and the future will show if new initiatives (e.g. the PDBdev [9]) may change that.

So far, intra- and inter-molecular movements [10,11] (herein collectively referred to as *protein dynamics*) are still challenging to detect experimentally, while precisely these protein dynamics determine if, for example, our muscles contract [12,13], our eyes can see [14], or our neurons conduct properly [15]. Herein, we focus on protein dynamics ranging from small loop motion to collective domain motions and complex rearrangements (**Fig. 1**), which occur on diverse timescales from sub-microseconds to many minutes [16,17], reflecting the broad range of energy barriers involved. Quantifying the timescales and order in which protein dynamics occur is the key to understanding these energies and molecular driving forces. Ideally, corresponding experiments would reveal the intricate interplay of dynamics on fast and slow timescales – i.e. low and high energy barrier crossings – which eventually lead to protein function. In reality, however, the accessible range of timescales is limited by the temporal bandwidth of the experiment at hand. In general, there are three ways to quantify biomolecular dynamics: (i) quantifying structural flexibility e.g. using root-mean-square deviations (RMSDs) in units of space rather than time; (ii) quantifying mean residence times in specific functional states; (iii) quantifying also forward and reverse rate constants between individual functional states A, B, C, etc. While all three provide valuable information, only the third approach can provide the kinetic and energetic information that is needed to identify which processes occur in thermal equilibrium and which represent out-of-equilibrium processes driven by an external energy source. In this article, we describe an extension to approach (iii) aimed at improving the detection of large-energy-barrier crossings in the ≥ms range, which are the decisive rate-limiting steps in protein function.

**Figure 1:**
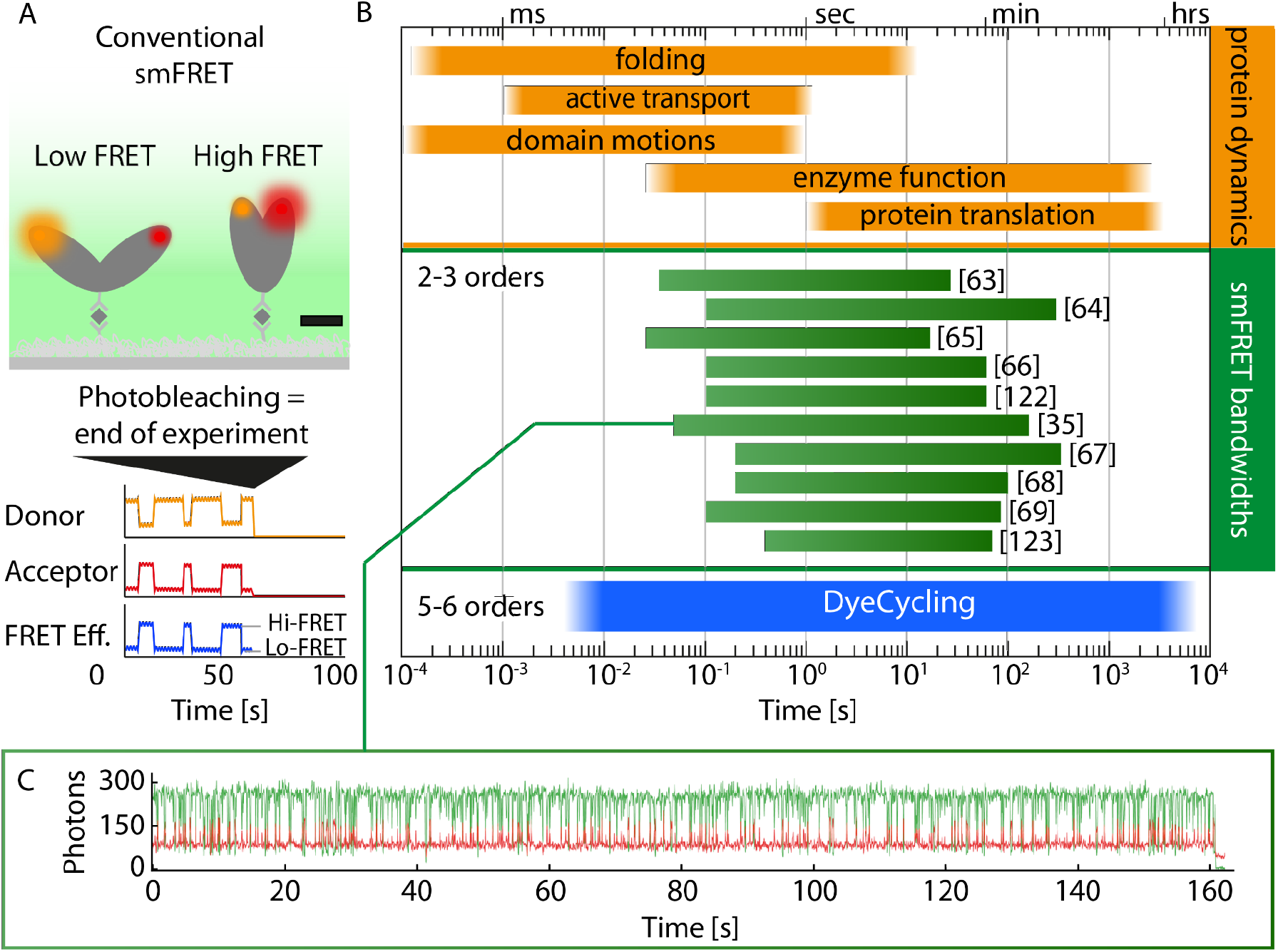
Timescales of protein dynamics and accessible bandwidths in conventional smFRET. ***(A)** Illustration of a conventional smFRET experiment with a surface-immobilized biomolecule (gray) with covalently attached donor (orange) and acceptor (red) dyes. Scale bar: 5nm. Irreversible photobleaching marks the untimely end of such smFRET recordings as indicated schematically. **(B)** The broad range of protein dynamics spanning >8 orders in time. Best case bandwidths from published smFRET studies (detailed in **Table 1**) cover only 2-3 orders in time. DyeCycling (blue bar) can double the accessible range to 5-6 orders in time. **(C)** smFRET trajectory with (to our knowledge) the broadest temporal bandwidth found in literature* [35]. *It was recorded in a semi-reversible binding scheme as discussed in the text*.

### 1.2 Single-molecule FRET can reveal the functional cycle of biomolecules

Individual state transitions of proteins or ribozymes are normally undetectable (averaged out) when ensembles of unsynchronized molecules are measured in bulk. Several single-molecule methods have been developed to reveal them nevertheless – one molecule at a time [18]. A very popular and versatile technique makes use of fluorescence resonance energy transfer to reveal the progression of a single biomolecule through its functional cycle. FRET can occur between two fluorophores within less than 10nm distance. The physical basis is a dipole-dipole interaction between a so-called FRET donor and a (usually red-shifted) FRET acceptor dye which scales with r^-6^, offering sub-nanometer distance resolution in smFRET experiments [2]. In short: the closer the two dyes are in space, the more efficient is the energy transfer from the excited donor to the acceptor dye, and the higher (lower) is therefore the observed acceptor (donor) intensity, respectively. As these fluorescence intensities can be conveniently detected at the single-molecule level, smFRET recordings became a powerful way to study the nanodynamics of biomolecules, which is widely used today throughout the life sciences [3,4]. A typical experiment to measure time-resolved smFRET trajectories (**Fig. 1A**) involves (i) site-specific labelling of the biomolecules of interest with a suitable FRET pair consisting of two organic dyes; (ii) the immobilization of the biomolecule on a passivated microscope slide, e.g. by biotin-avidin coupling; (iii) the recording of spectrally split single-molecule fluorescence trajectories, e.g. using total-internal reflection (TIR) excitation and highly parallelized camera-based detection, or alternatively using confocal detection and avalanche photodiodes. In this way, single-molecule dynamics can be observed in real time as fluorescence-intensity changes reflecting the inter-dye distance changes in the biomolecule.

## 2 State of the art & current limitations of smFRET

### 2.1 Photobleaching – the fundamental limit of conventional smFRET

SmFRET studies are fundamentally limited by irreversible photobleaching of one of the two dyes involved, which terminates the observation of a given biomolecule [19]. As shown in **Figure 1B**, the currently achieved temporal bandwidth of smFRET experiments (defined as the observation time divided by the time resolution of the experiment, Δ*T*/Δ*t*) spans only 2-3 orders of magnitude, and it has not markedly improved in the past ten years (see **Table 1**). (We note that the displayed values represent best-case bandwidths based on the longest, most informative trajectories depicted in scientific publications, while most recorded trajectories are much shorter due to the exponentially distributed trajectory lengths caused by stochastic photobleaching.) Ergo, at present, only hundreds to a few thousand datapoints can be measured before irreversible photobleaching occurs. This statement holds on diverse timescales, because faster recordings require higher laser power to achieve a useful signal-to-noise ratio, which inevitably causes faster photobleaching [20]. The smFRET detector is thereby not the limiting factor: the quantum yield of current detectors exceeds 90%, and technically single photons can be detected with a resolution of picoseconds, using avalanche photodiodes [21]. Still, most time-resolved smFRET trajectory studies use highly parallelized (EMCCD or s-CMOS) camera detection offering down to millisecond time resolution; while at a useful signal-to-noise ratio, smFRET trajectories recorded with 1 ms resolution last for 2-3 seconds before photobleaching [22]. So rather than the detector, the limited photostability of the fluorescent probes is the main cause of the narrow bandwidths in current smFRET studies. Aiming for as small and as photostable probes as possible, small organic fluorophores are the preferred choice for precise time-resolved smFRET detection. Unfortunately, despite intense research [19,23–26], the bleach rate of dyes used in smFRET studies has seen little improvement over the past ten years (cf. **Table 1**), and ATTO 647N is still the most photostable dye at present [27].

**Table 1:** Best case literature values of time resolution, observation time, and resulting bandwidth in surface-immobilized smFRET. ALEX: Alternating Laser EXcitation.

| Time resolution<br>$\Delta t$ [ms] | Observation time<br>$\Delta T$ [s] | Bandwidth<br>$\Delta T / \Delta t$ | Sample | Reference |
| --- | --- | --- | --- | --- |
| 33 | 27 | $8.1 \times 10^2$ | RNA | [63] |
| 100 | 255 | $2.5 \times 10^3$ | Protein-DNA | [64] |
| 25 | 18 | $7.0 \times 10^2$ | Protein-RNA | [65] |
| 100 | 60 | $6.0 \times 10^2$ | Protein | [66] |
| 100 | 60 | $6.0 \times 10^2$ | Protein-DNA | [122] |
| 50 | 158 | $3.1 \times 10^3$ | Protein-protein | [35] |
| 200 | 270 | $1.3 \times 10^3$ | Protein-RNA | [67] |
| (ALEX) 200 | 85 | $4.2 \times 10^2$ | Protein | [68] |
| (ALEX) 100 | 25 | $2.5 \times 10^2$ | Protein | [69] |
| (ALEX) 400 | 70 | $1.7 \times 10^2$ | Protein-DNA | [123] |

In view of the broad-range dynamics of proteins (**Fig. 1B**), an accessible time window of 2-3 orders of magnitude is very limiting. For example, correlations between fast (ms) and slow (min) dynamics have remained inaccessible by smFRET, since they are lost when separate datasets are measured at fast and slow sampling rates. Moreover, even the initial trajectory selection (distinguishing meaningful trajectories from experimental artefacts) can be ambiguous if not all of the facets of a biomolecule’s behavior occur within the short photobleaching-limited observation time. Also, static disorder (i.e. lasting differences in the behavior of individual molecules, or subpopulations of the ensemble) can hardly be revealed using the current narrow bandwidth. This matters, because almost all smFRET studies, rely on the ergodic assumption which states that the time average and the ensemble average of a given biomolecular system are identical [28,29]. In other words, this means that at least theoretically, the full ensemble behavior of a given biomolecule can be recovered from the – very long – observation of just one single molecule. Apparent ergodicity breaking in biomolecules has been described for protein [30], ribozyme [31–34], and DNA systems [29] alike. It can result, for example, from complex energy landscapes where different conformations are not or only rarely interconvertible (limited by the available energy sources). However, since ergodic equilibration times can take very long for large biomolecules with rugged energy surfaces [29], the fundamental ergodic assumption can often not be confirmed in smFRET studies.

In this context, it is worth taking a closer look at the experimental implementation of the largest bandwidth in **Fig. 1C** by Zosel *et al*. [35]. It was achieved using a biological system that shows transient proteinprotein interactions: the protein labelled with the Cy3B donor dye was surface-immobilized, while the protein labelled with the acceptor dye ATTO 647N underwent reversible binding and release. Consequently, this experiment is only limited by donor (but not acceptor) photobleaching. In addition, confocal detection was used, and photobleaching was reduced by use of an argon-saturated buffer as well as an oxygen scavenger system. While, even under such carefully optimized measurement conditions, the three orders of magnitude in time couldn’t be surpassed, this landmark data hints at an interesting way forward for surface-immobilized smFRET studies of biomolecular dynamics.

### 2.2 The reversible dye binding trick

Reversible dye binding is a promising way to decouple the smFRET observation time from the currently prohibitive dye bleaching. While so far ignored in time-resolved smFRET studies, this trick is routinely used in super-resolution imaging, e.g. in point accumulation in nanoscale topography (PAINT) described in 2006 [36]. Later, oligonucleotide hybridization appeared as a convenient way to tune dye-binding kinetics leading to DNA-PAINT [37]. FRET entered the stage as FRET-PAINT which could accelerate imaging rates by suppressing the background signal at higher dye concentrations that enabled faster binding rates [38]. Beyond super-resolution imaging, reversible oligonucleotide-based dye binding was found to be useful for the sequential detection of multiple FRET pairs within one single molecule, a technique termed FRET X, that was developed to identify proteins based on characteristic FRET fingerprints [39]. While all these applications disregard the time domain information, recently a first time-resolved study showed that particle tracking can profit greatly from reversible dye binding [40]. By contrast, time-resolved smFRET experiments still adhere to static covalent dye labelling, where a single photobleaching event puts an irreversible end to the experiment resulting in narrow temporal bandwidths throughout the field (**Table 1**). To overcome this prohibitive limitation, we introduce here the concept of DyeCycling for the study of biomolecular dynamics in the millisecond to hour range. Standing on the shoulders of giants, DyeCycling expands the strategy of reversible dye binding into a versatile labelling scheme, aimed to vastly expand the time range covered by smFRET trajectories.

## 3 Breaking the photobleaching limit in time-resolved smFRET with DyeCycling

### 3.1 The DyeCycling concept

If dyes bleach, why don’t we replace them with new ones? DyeCycling makes use of reversible dye binding to the biomolecule of interest to decouple the total observation time from photobleaching (**Figure 2A**). Several (bio-)chemical strategies enable such reversible dye binding with tunable kinetics (**Section 3.2** and **Figure 3**). Background fluorescence of unbound dyes is suppressed by localized excitation and detection schemes, such as total internal reflection (TIR), confocal detection, or using zero-mode waveguides (**Section 3.3**). Site-specific binding and dissociation of donor and acceptor dyes happens spontaneously and continuously, leading to the – in theory – endless observation of the dynamic behavior of a surface-immobilized biomolecule via smFRET recordings. The resulting smFRET trajectory is comprised of four different phases: the biomolecule is either in the ‘FRET regime’ with acceptor and donor bound, in the ‘donor only’ or ‘acceptor only’ regime where just one dye is bound, or in the ‘dark regime’ without any functional dyes present (illustrated in **Fig. 2B** left). The goal is now to maximize the residence time in the FRET regime, which can be achieved by tuning the dye binding and dissociation rates, plus the sampling rate of the measurement – all while considering the dye-specific bleach rate. A Monte Carlo simulation based on experiment-derived rate constants [37,41–43] illustrates the situation (**Fig. 2C**). It shows the DyeCycling-facilitated hour-long detection of a hypothetical biomolecule alternating between three functional states as illustrated by the kinetic 3-state model in **Fig. 2B** (right). The zoomed-in view (**Fig. 2C** bottom) shows that a conventional smFRET experiment would not be able to resolve how this biomolecule proceeds through its fast and slow dynamics: either the static low-FRET pieces would be sorted out as non-functional artefacts, or slower measurement settings would be chosen leading to time averaged mixing of the states S1 and S2. In contrast, the DyeCycling trajectory connects the short pieces into one sequential single-molecule observation, offering conceptually new information about the biomolecule’s broad-range dynamics. The (experiment-derived) simulation resulted in 80% FRET regime during which the biomolecular dynamics are resolved. The remaining gaps can be dealt with during analysis, e.g., using Hidden Markov models (implemented in SMACKS [44] or other software packages [45]) to describe the biomolecular states, and additionally three states for the ‘dark’, ‘donor-only’, and ‘acceptor only’ regimes. In this way, the quantization of the observed biomolecular kinetics is not affected by the intermittent pauses. Experimentally, DyeCycling can run autonomously on one (focus-stabilized) field-of-view with no user input needed during the hourlong recordings, and post-hoc x-y-drift correction allows the extraction of hour-long fluorescence trajectories as shown in **Section 4**.

**Figure 2:**
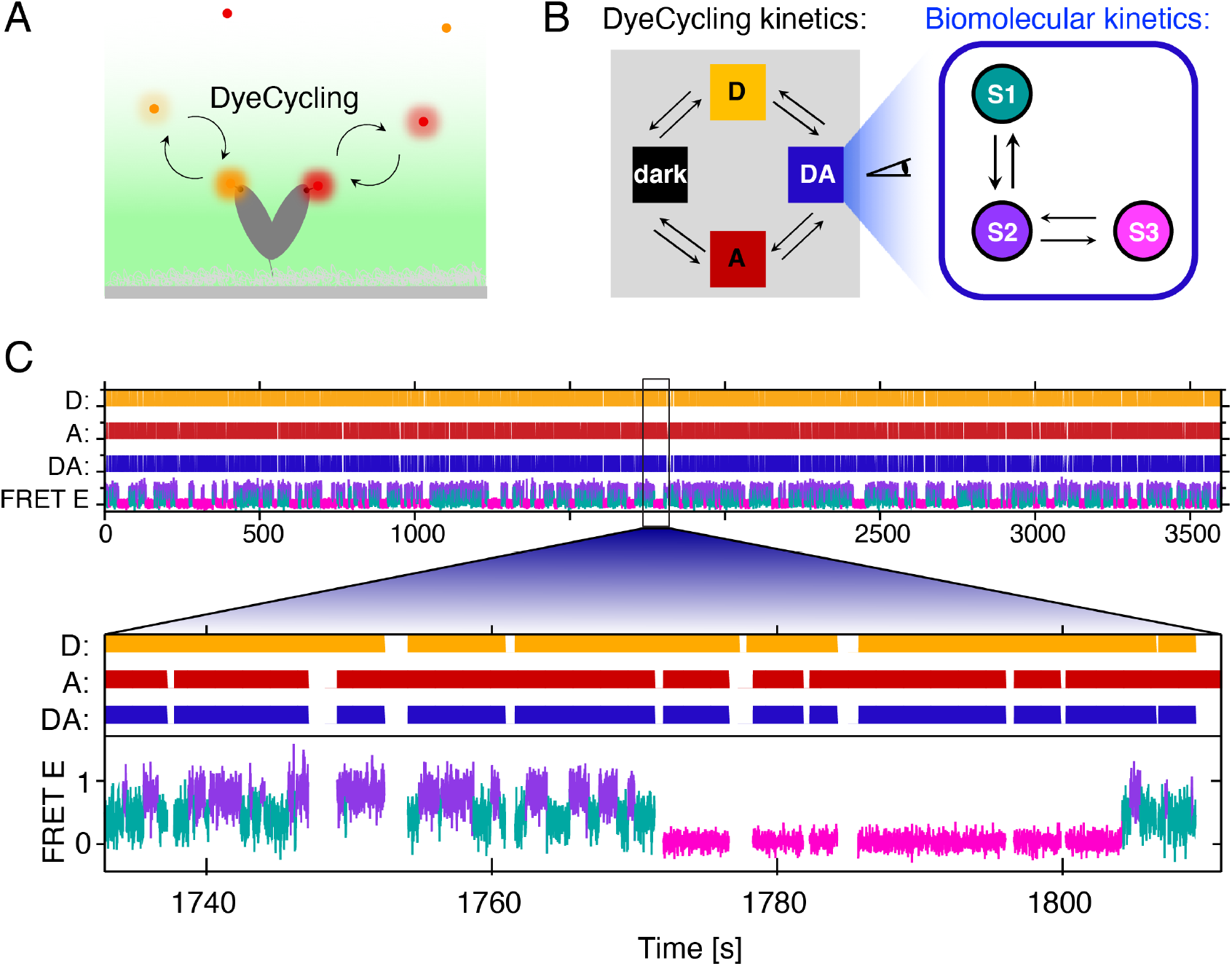
DyeCycling to detect the dynamics of a single molecule from milliseconds to the hour range. **(A)** Illustration of a DyeCycling experiment with an immobilized protein, reversibly labelled by ‘dye cyclers’ (donor cycler: orange, acceptor cycler: red). **(B)** Illustration of the kinetic models used in the Monte Carlo simulation of DyeCycling. The DyeCycling kinetics (reversible dye binding) was simulated using a 4-state model: no dye bound: ‘dark’, only donor bound: ‘D’, only acceptor bound: ‘A’, donor and acceptor bound: ‘DA’, time resolution: 10ms, D or A binding rate: 1s^-1^, D or A dissociation rate: 0.1s^-1^. The biomolecular kinetics were simulated using a 3-state model with fast (k_12_ = k_21_ = 1s^-1^) and slow (k_23_ = k_32_ = 0.1s^-1^) dynamics. **(C)** Resulting simulated smFRET trajectory with reversible donor and acceptor binding leading to 80% co-labelled FRET regime with intermittent pauses. The FRET efficiencies (FRET E) and Gaussian noise (standard deviations) of the three biomolecular states were set to: E_1_ = 0.4 ± 0.2, E_2_ = 0.8 ± 0.2, E_3_ = 0.05 ± 0.1. Color code as in (B).

**Figure 3:**
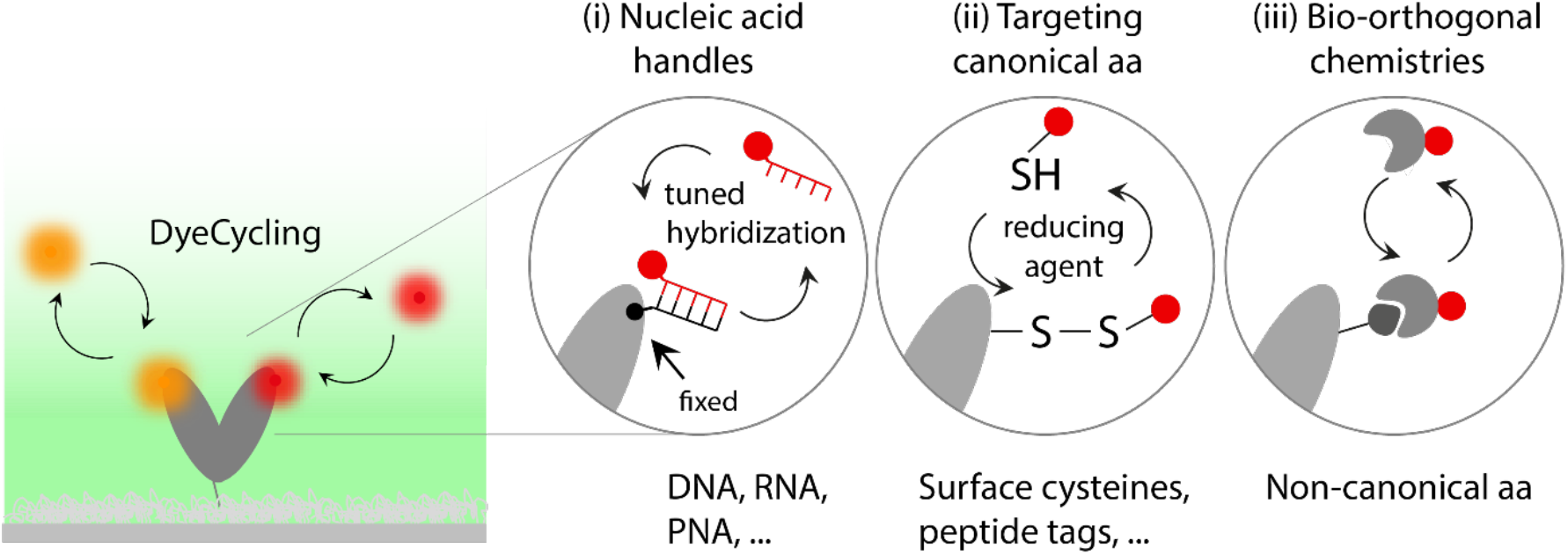
Reversible dye labelling strategies for DyeCycling. (**i**) Nucleic acid handles (DNA, RNA, PNA, etc.) covalently attached to the protein of interest accommodate dye-labelled complementary cycler strands. (**ii**) Several canonical amino acids allow for site-specific reversible binding, e.g. cysteines and thiolated dyes. (**iii**) Non-canonical amino acids give access to diverse bio-orthogonal chemistries to introduce reversible dye binding.

Altogether, DyeCycling has the potential to vastly expand the time range covered by smFRET trajectories, enabling the intermittent observation of a single biomolecule for an hour with a time resolution of milliseconds, i.e. covering 5-6 orders of magnitude in time. Compared to conventional smFRET with covalently attached dyes, this can provide 100-1000-fold more measured datapoints – i.e. *more information* – per single molecule. Consequently, data reliability and trajectory selection will improve by the same factor. Most importantly however, new dynamic effects, such as correlations between fast and slow dynamics, ergodicity breaking, etc. may become accessible.

### 3.2 Cycler probe designs

Site-specific reversible dye binding in DyeCycling can be achieved with various chemistries and cycler designs. DNA handles (**Fig. 3i**) are an obvious choice borrowed from super-resolution imaging [37]: an ssDNA oligo may be covalently attached to a protein of interest as a docking target for a complementary dye-cycler oligo. A wealth of information is available on hybridization kinetics [37,41,43,46]. The hybridization rate depends on the cycler’s diffusion time, and is tunable with cycler concentration, temperature, and salt concentration, whereas the dissociation rate can be tuned via the DNA duplex length and sequence (GC content, mismatches [41,47]), salt conditions, and temperature [41]. In addition, synthetic nucleic acid analogues have been developed to overcome potential shortcomings of DNA [48], such as its negative charge, relatively low affinity, and susceptibility to degradation. One such analogue is peptide nucleic acid (PNA), which consists of a pseudopeptide backbone (*N*-(*2*-aminoethyl)glycine), with nucleobase side chains [49] that undergo Watson-Crick base pairing like regular DNA, but with much higher affinity which enables smaller probes (~5bp) [50] with faster diffusion-limited on-rate. Dye labelled PNA probes have been successfully implemented as erasable dyes in livecell imaging [51], and custom-functionalized PNA is commercially available [52,53].

Alternatively, surface cysteines may be used for reversible disulfide bridge formation under reducing conditions, to circumvent the need for artificial docking targets (**Fig. 3ii**). Single-molecule force spectroscopy and optoplasmonic studies showed thiol-disulfide reversibility with kinetics that are suitable for reversible site-specific dye labelling [54,55]. This direct labelling concept can be extended beyond cysteines to other (rare) canonical amino acids or small peptide tags that can be introduced by site-specific mutagenesis. A well-known example is the poly-histidine tag that binds to nickel-nitrilotriacetic acid (Ni-NTA). Several other peptide tags and different metal ions have been studied in the context of specific protein labelling [56]. Since metal chelation is non-covalent, it lends itself for reversible binding applications where the binding affinity can be varied, e.g. by the number and arrangement of histidines in a tag [57]. Additionally, unnatural amino acids offer further options to tune metal chelation kinetics [58]. Indeed, the development of such non-canonical amino acid incorporation by genetic code expansion [59,60] (**Fig. 3iii**) opens near endless possibilities to site-specifically introduce a desired bio-orthogonal binding capacity. While, traditionally, the highest stability and irreversible coupling was usually pursued [61], DyeCycling asks for the opposite: reversible binding with specific kinetics. Ideally, short dye linkers could be introduced to improve the distance resolution in FRET space [62].

The main requirements for optimal cycler probes are the same in all three cases: fast and site-specific cycler binding, photostable dyes with a slow bleach rate, and a cycler dissociation rate that is optimized accordingly. Using a literature example of a dye binding rate constant of k_on_ = 6×10^6^ M^-1^ s^-1^ [42] and a cycler concentration of 150 nM, yields a binding rate of r_on_ = 0.9 s^-1^, ca. 1 binding event per second. Thus, a dissociation rate of r_off_ 0.1 s ^-1^ leads to a bound fraction of 90%, and to average bound times of 10s of the individual cycler. Good, photo-stable organic dyes last for an average of 1000 datapoints under optimal experimental conditions, thus 10ms is a suitable time resolution in this case (ie. 100 Hz sampling rate). Useful dye candidates are (amongst others) Cy3, Cy3B, Cy5, LD550, LD655, ATTO 550, and ATTO 647N [35,63–69], as they show stable emission rates, high quantum yields, and relatively large photon budgets, while ultimately, the optimal dye choice may differ per biomolecule and local dye environment.

### 3.3 Background suppression by localized excitation and detection schemes

With many fluorescent cycler probes present in solution, DyeCycling requires strong suppression of fluorescent background. Total internal reflection fluorescence (TIRF) microscopy is a simple and widespread technique to establish localized excitation at the glass-water interface of the sample chamber, thereby reducing background excitation and thus enabling the detection of single fluorophores [70]. As demonstrated below, TIRF provides sufficient background suppression for the resolution of individual biomolecules under DyeCycling conditions, which makes DyeCycling readily available to many smFRET experimentalists. Moreover, thanks to the parallelized widefield detection of hundreds of individual biomolecules at once, TIRF combined with DyeCycling can offer large amounts of information in just one measurement. Alternatively, confocal detection using avalanche photo diodes offers single-photon counting [35], but the gained ultimate time resolution comes at the cost of detecting just one molecule at a time. In contrast, zero-mode waveguides (ZMW) offer much more localized excitation combined with parallelized recordings in arrays, as used e.g. in commercialized PacBio DNA sequencers [71]. Such metal nano-wells are particularly interesting for DyeCycling because with their zeptoliter observation volumes, they enable single-molecule detection at 1000-fold higher dye concentrations (~100μM [72]). This facilitates faster dye binding rates which can make even shorter timescales accessible in DyeCycling experiments. The newest palladium-based ZMW generation is chemically inert and autofluorescence free [73]. In addition, multi-color detection (>2 colors) could be implemented e.g. using prisms [74,75] to spectrally split individual colors in-situ, without the need for multiple separate detection channels. In summary, TIRF is sufficient to implement DyeCycling as demonstrated below, and zero-mode waveguides can further improve DyeCycling and extend it towards short timescales.

## 4 Experimental proof-of-concept of DyeCycling

### 4.1 The hour-long observation of cycling dyes

As a biomolecular test system for DyeCycling, we chose a Holliday junction [29,76], i.e. a four-way DNA junction found in homologous DNA recombination. Our Holliday junction design (**Fig. 4A, Table S1**) features two DNA overhangs to accommodate two dye-labelled DNA oligos, termed donor and acceptor cyclers. **Fig. 4B** shows the workflow of a DyeCycling experiment: as expected, the surface immobilization of the unlabelled Holliday junction leaves a spotless field of view as a starting point (Fig. 4B i). The Holliday junction is then labelled in-situ by adding the donor cyclers to the measurement buffer, which hybridize site-specifically to the complementary Holliday junction overhang, resulting in bright spots in the donor channel only (Fig. 4B ii). Unbound cyclers are not resolved as spots (at 60ms exposure time) because they diffuse too fast, and under TIRF illumination, they only moderately increase the background fluorescence at 100nM (Fig. 4B iii). When also acceptor cyclers are added to the buffer, they hybridize to their complementary Holliday junction overhangs, leading to bright spots in the acceptor channel and in the FRET channel (Fig. 4B iv). The latter indicates an inter-dye distance of <10 nm (since direct excitation of the acceptor is negligible at 520nm), which denotes the simultaneous binding of both cyclers to one Holliday junction molecule. Specific reversible labelling can easily be distinguished from rare non-specific adsorption given the hour-long DyeCycling observations (exemplified in **Fig. 4C**), which unambiguously identify the positions of the reversibly labelled Holliday junctions. Such long trajectories could be recorded for more than fifty biomolecules in parallel (with 256×512 pixels), each showing continuous binding and dissociation (or bleaching) events of the donor and acceptor cyclers. The associated histograms show a cycler-bound coverage of just 50% in time, which may still be improved given that binding rates of 1/s have been achieved in similar experiments [37] (see also next section).

**Figure 4.**
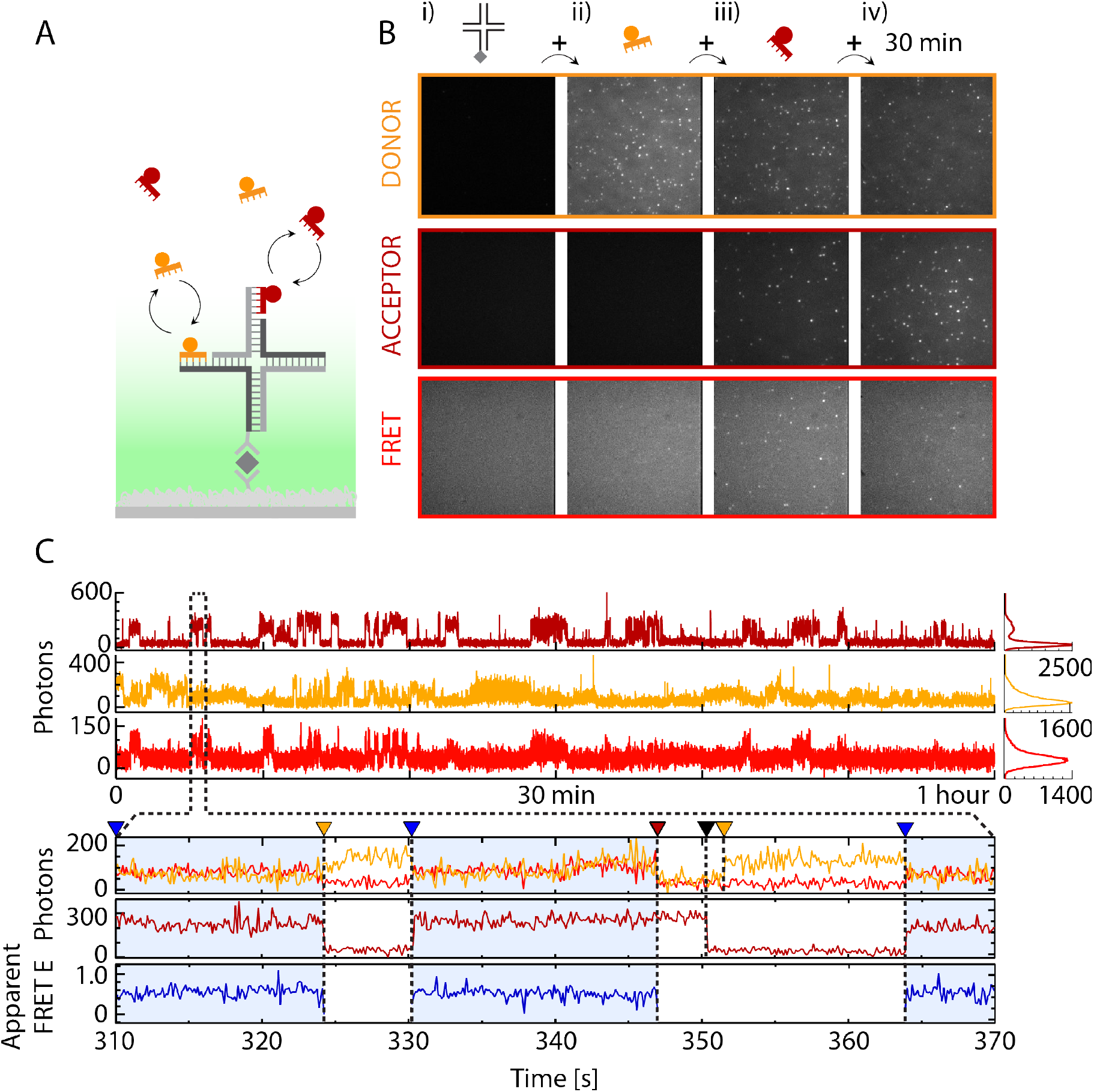
Hour-long DyeCycling on immobilized Holliday junctions. (**A**) Schematic of a surface-immobilized Holliday junction with donor (orange) and acceptor cyclers (dark red). Oligo sequences are found in **Supplementary Table S1**. (**B**) Workflow of a DyeCycling experiment shown by sequential fields of view of the donor channel (donor emission after donor excitation), the FRET channel (acceptor emission after donor excitation), and the acceptor channel (acceptor emission after acceptor excitation). 100 nM cyclers were recorded with 60 ms exposure time (see Methods in **Section 4.3**). Donor and acceptor channel: 25-frame average, FRET channel: 50-frame averages, each with identical contrast per channel. (**C**) Corresponding hour-long fluorescence trajectory of a single Holliday junction with repeated cycler binding and dissociation. The zoom view shows a binding event of both cyclers, resulting in FRET (blue-shaded area). Triangles show (left to right) the start in the FRET-regime (blue), acceptor cycler leaving (orange), acceptor cycler arrival leading to FRET (blue), donor cycler leaving (dark red), acceptor cycler leaving (black), donor cycler arrival (orange), acceptor cycler arrival leading to FRET (blue).

The zoom view in **Fig. 4C** shows the progression of a single Holliday junction from an initial FRET-regime (blue shading), to a donor-only regime caused by acceptor-cycler dissociation or bleaching (orange triangle). Shortly after, a fresh acceptor cycler binds (blue triangle), and the resulting FRET-regime lasts until bleaching or dissociation of the donor (dark red triangle) and later the acceptor cycler (black triangle). Next, another donor cycler binds (orange triangle) followed by an acceptor cycler (blue triangle) initiating the next FRET-regime etc. The donor trace shows generally a higher noise level compared to the acceptor trace, which can only partly be attributed to quenching by FRET. In the future, this may be improved by more advanced excitation schemes, such as zero-mode waveguides, to further reduce background fluorescence. While, with this first prototype of a DyeCycling construct and partially optimized conditions, conformational dynamics were hardly detectable (as discussed below), in comparison to conventional smFRET with covalent labelling, we gained an encouraging factor of twenty in doubly-labelled observation time thanks to DyeCycling (as estimated from Gaussian histogram fitting). We conclude that further measures are needed to improve the signal-to-noise ratio, but that the near endless cycling of dyes is feasible, as demonstrated by the repeated cycler binding and dissociation events observed on one single biomolecule for up to an hour.

### 4.2 Current shortcomings, solutions, and fundamental limits

Based on this proof-of-concept study, we identify several points to improve that fall into two categories: setup- and construct-related aspects. Most of the current experimental limitations can be solved in the future. On the setup side, the time resolution (2×60ms ALEX) was limiting for the fast Holliday junction dynamics at hand – even using conventional smFRET with covalently attached dyes (**Supplementary Fig. S1**). The limiting factor was the green laser power available (rather than the detector time resolution of milliseconds), which limited the photon emission rate. In addition, the finite background-suppression by TIRF limited the feasible cycler concentration (100 nM) and therefore the cycler binding rate, which in turn limits the time resolution of the experiment and the fraction of time spent in the FRET-regime (donor- and acceptor-cycler bound). Zero-mode waveguides may solve this issue in the future, since they offer a much-reduced observation volume (zeptoliter [77]) which allows for higher cycler concentrations (tens to several hundreds of μM [72]). On the construct side, the cycler design can be further optimized, e.g. to tune the dissociation rates via the hybridizing DNA sequence, or using shorter PNA instead. Moreover, some Holliday junction constructs showed substantial collisional quenching, indicating that dye attachment was suboptimal. The current dye-DNA attachment resulted in a long linker (ca. 1.5 nm contour length) which can increase fluorescence quenching by dye-DNA interactions [78]. Naturally, more photo-stable dyes with a smaller bleach rate would benefit DyeCycling as well as conventional smFRET [26].

Encouragingly, non-specific cycler adsorption (sticking) caused no problems using standard polyethylen glycol (PEG) passivation, even above 100nM cycler concentrations. This could become more challenging using protein samples, which we plan to solve using multi-PEG passivation or lipid bilayer passivation [79,80]. Vertical drift – during the hour-long measurement – was efficiently suppressed by feedback-controlled focus stabilization, and horizontal drift could be easily corrected post-hoc, using in silico drift correction (tens to hundreds of nanometers per hour). Also, the datafile size is manageable, with 32.7 GB for a one-hour measurement of 512×512pixels at 60 ms per frame. Lastly, beyond the experimental shortcomings that can and will be addressed in future work, DyeCycling is limited by two fundamental effects: the time resolution is limited to milliseconds due to the unwanted detection of diffusing molecules at even shorter timescales [81], and the total observation time is ultimately limited by the stability of the surface-immobilized biomolecule [82]. So far, we demonstrated the detection of a biomolecule, reversibly labelled with a dye-cycler FRET pair, over the duration of an hour with 120ms time resolution.

### 4.3 Methods

A published Holliday junction [83] was modified with two overhangs to accommodate hybridization of a donor and an acceptor cycler oligo (**Supplementary Table S1**). HPLC-purified oligo strands (Biomers, Germany) were diluted to 1 μM in TN50 buffer (10 mM Tris-HCl, 50 mM NaCl, pH 8; filtered with a Corning Syringe filter with a 0.22 μm nylon membrane (Merck, Germany)), and annealed with a thermocycler (T-Gradient Thermoblock, Biometra, Germany) for 10 minutes at 90°C followed by cooling to 20°C (1°C/min). PEGylated slides for single-molecule measurements were prepared as previously described [84]. In short, slides were burned (90°C/h heating up, 1h at 500°C, 150°C/h cooling rate), functionalized with Vectabond (3-aminosilane, Vector Labs, USA), and passivated with a mixture of 20% (w/v) polyethylene glycol (mPEG-succinimidyl valerate MW 5000, Laysan Bio Inc., USA) and 0.75% (w/v) biotinylated PEG (biotin-PEG-succinimidyl carbonate MW 5000, Laysan Bio Inc., USA) in MOPS buffer (50 mM 4-morpholinepropane sulfonic acid, pH 7.5, 0.22 μm-filtered). Measurements were performed in silicone culture well gaskets (Grace Biolabs, USA) stuck onto the functionalized slide. The measurement buffer consisted of TN50, 1% gloxy (1 mg/ml glucose oxidase (Sigma, USA) plus 0.04 mg/ml catalase (Roche Diagnostics, Switzerland)), 1% (w/v) glucose, and 1 mM Trolox for oxygen scavenging and triplet state quenching [85]. Sample and measurement buffer were mixed right before the measurement. All buffer chemicals were purchased from Sigma-Aldrich.

Single-molecule measurements were performed on a previously described TIRF setup [86] with an EMCCD camera (iXonUltra 897, Andor, United Kingdom) and autofocus stabilization (MS-2000, Applied Scientific Imaging, USA), using alternating laser excitation (ALEX) [87] with 60 ms exposure time (i.e. ALEX time resolution of 120 ms) with fiber-coupled diode lasers (520 nm and 638 nm, Lasertack, Germany). Measurements were performed at 13 mW (green) and 5.5 mW (red), measured directly after the fiber. The sample well was incubated with 20 μL neutravidin (0.25 mg/mL, ThermoFisher, USA) for 5 min, washed with 600 μL TN50 buffer, incubated with 20 μL unlabelled Holliday junction (10 pM in TN50 buffer) for 1 min, and washed with 600 μL TN50 buffer. Donor cycler and acceptor cycler were diluted in measurement buffer to 100 nM, 20 μL were used for the measurement, and the well was covered with a coverslip. Acquired movies were corrected for horizontal drift using the *Estimate drift* functionality of the ImageJ NanoJ Core plugin [88]. Data were analyzed in Igor Pro (Wavemetrics, USA) using existing code developed in the lab of Thorsten Hugel, and new code to extract DyeCycling trajectories from multiple movies.

## 5 Biological use cases

DyeCycling can be of widespread utility, since it is a general feature of biomolecules that slow functional dynamics are based on a hierarchy of faster dynamics. Moreover, the study of slow ergodic relaxations requires single-molecule observations that cover many orders of magnitude in time. Out of nature’s vast biomolecular pool, we highlight here a selection of vital protein systems (**Fig. 5**) which illustrate the wide span of applications of DyeCycling, to elucidate how the interplay of fast and slow dynamics causes protein function.

**Figure 5:**
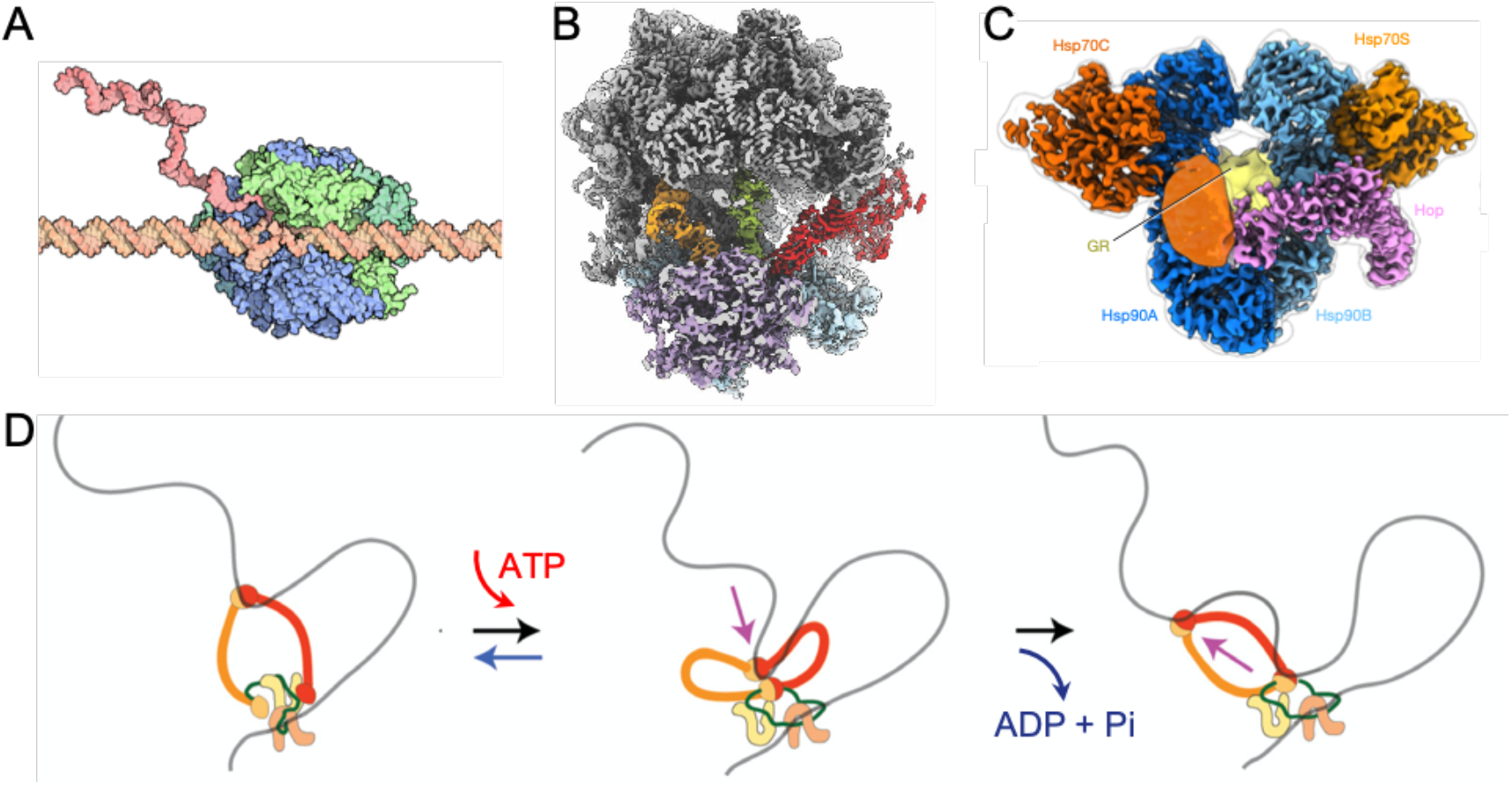
Use cases for DyeCycling. *Four essential protein systems that are rich in broadrange dynamics and may therefore benefit from the width of accessible timescales offered by DyeCycling. A) Transcription by RNA polymerase (pdb:1i6h)* [120]. *B) Translation by the ribosome* [121]. *C) Chaperone action by Hsp90* [110]. *D) DNA loop extrusion by the SMC protein condensin* [89].

### 5.1 Transcription and translation machineries

The central dogma of molecular biology – DNA is transcribed into RNA, RNA is translated into proteins – is rife with dynamic heterogeneity [89]. Pauses during RNA polymerase activity have been shown to be essential in regulating transcription and thus gene expression [90–94]. Using magnetic tweezers, pausing events of bacterial RNA polymerase could be studied with a high temporal bandwidth (2.5h with 40ms resolution, i.e. 5 orders in time), which revealed considerable heterogeneity in transcription velocity and pause dynamics [95]. While the causative internal rearrangements of the polymerase remain elusive in the magnetic tweezers assay, they could be resolved in real-time smFRET recordings [92], albeit at a much-reduced temporal bandwidth. DyeCycling may fill this (bandwidth) gap by bridging from milliseconds to the hour range, and reveal new disease-relevant insights on pausing heterogeneity, as well as, static (molecule-to-molecule) heterogeneity.

SmFRET has also been used to study protein translation, e.g., to reveal t-RNA fluctuations [96], proof reading mechanisms [97], or competing interactions [98]. However, these studies are similarly limited by a total observation time of seconds [97], while protein synthesis extends towards the hour range with an approximate speed of 5-15 amino-acids per second [99,100]. Again, this illustrates the necessity of an increased width of accessible timescales in smFRET to meet the broad timescales involved in biomolecular function.

### 5.2 Hsp90 ‘chaperones’ the proteome

The molecular chaperone heat-shock protein 90 (Hsp90) is a central regulator of proteostasis [101] that is involved in the function of 20% of the proteome [102]. For this purpose, Hsp90 undergoes a plethora of transient interactions [103] with assisting cochaperones and diverse client proteins including regulatory kinases, hormone receptors, and transcription factors, such as the tumor suppressor p53, making it a central drug target for anti-cancer therapy [104,105]. Despite its biomedical importance and the related intense research activity over the past decades [106–108], even fundamental functional aspects have remained enigmatic, such as the role of Hsp90’s notoriously slow ATPase function for the chaperoning action [109]. Two recent cryoEM structures provide an unprecedent molecular view on the intricate multipartite complexes involved in client loading and maturation of the steroid hormone receptor GR by Hsp90 [110,111], however the dynamic time-domain information on client processing lags once more behind its 3D structural counterpart. While bulk assays have revealed client refolding rates on the order of many minutes [112,113], and Hsp90 undergoes characteristic large conformational opening-closing transitions in the range of milliseconds to minutes [114], it has remained challenging to link client processing with precise conformational dynamics in the Hsp90 system – partly due to the broad range of timescales involved. By increasing the smFRET-accessible time range with DyeCycling, we expect to capture also transient (ie. short and rare) events with high temporal resolution and to set them into the broad timescale perspective, in order to elucidate the active side of Hsp90’s chaperoning action.

### 5.3 Structural maintenance of chromosome (SMC) proteins

SMC proteins, such as cohesin or condensin, are in charge of chromosome organization throughout all kingdoms of life, where amongst others, they play a key role in gene regulation and assist in proper chromosome segregation during cell division [115,116]. Single-molecule techniques show that, *in vitro*, they extrude large DNA loops (ten thousands of basepairs, i.e. several microns) in an ATP-dependent way, which could be observed in real-time and one loop at a time [117,118]. However, the molecular details of this active mechanical process remain more challenging to reveal. Recent findings [69,119] on cohesin hint now at a Brownian ratchet mechanism, where flexible large-scale (50nm) rearrangements occur in thermal equilibrium, while ATP binding and hydrolysis bias the system towards a directional sequence of events that causes processive loop extrusion. If true, this requires that ATP binding and hydrolysis modulate the individual binding affinity and/or rigidity of critical interaction sites and domains, but how such energy coupling occurs in cohesin remains to be uncovered. The existing smFRET data [69] supports this general concept, but the accessible temporal bandwidth of conventional smFRET limited the observation times to ≤20s with 50ms time resolution. DyeCycling could expand the total accessible timescales to capture more ATP hydrolysis cycles with sufficient time resolution, and to cover the loop-extrusion intervals for many minutes.

Clearly, there are many other protein systems with rich dynamics, e.g. those involved in DNA repair, protein degradation, protein secretion, and many motors, enzymes, etc., where often detailed snapshots of 3D structures are available. This allows researchers to take now the next step and address the underlying dynamics and energetics that ultimately cause protein function.

## 6 Summary

In this conceptual study, we introduced DyeCycling to break the photo-bleaching limit in single-molecule FRET. The reversible dye-binding scheme enables repeated, spontaneous dye replacements, and thus, the total observation time of a biomolecule is no longer limited by photobleaching of just a single dye (as in conventional smFRET). Instead, DyeCycling can vastly increase the number of data points – and therefore the information content – of time-resolved smFRET trajectories by a factor of 100-1000. Our first cycler prototypes yielded 20-fold more FRET datapoints per single molecule, albeit still at a low signal-to-noise ratio. Therefore, DyeCycling will be further developed to bridge from the fast millisecond timescales up to the minutes and hour range. Firstly, this allows to reach the slow timescales needed to investigate ergodicity breaking in complex biomolecules, such as proteins and ribozymes. Secondly, aiming to cover up to 6 orders in time, DyeCycling has the potential to reveal a wealth of new dynamic effects in biomolecules. As highlighted for a few protein systems, the interplay between fast and slow dynamics (i.e. low- and high-energy-barrier crossings) are fundamental to protein function, and DyeCycling can make them experimentally accessible. Thirdly, the significant increase in information per single molecule can improve the reliability of smFRET, since artefacts are more easily identified during the long DyeCycling trajectories. Fourthly, DyeCycling benefits the experimenter, since in contrast to conventional smFRET, this experiment can technically run for hours and thus provide a comprehensive dataset without the need of user input every few minutes. Further optimization of the setup components and the reversible binding kinetics is still needed to improve the experimental results. The direct extraction of the biomolecular kinetics can be implemented using machine learning tools, such as SMACKS [44]. In the future, we aim to apply DyeCycling to proteins, ribozymes, and other biomolecular systems, where the newly gained 5-6 orders in time and the >100-fold more information per single molecule suggests access to so far unexplored dynamic effects. Given its broad application range, we anticipate that DyeCycling will become a useful method to decipher the rich nano-dynamics of biomolecular systems.

## Acknowledgements

We thank Chirlmin Joo, Johannes Hohlbein, Mattia Fontana, and Abbas Jabermoradi for helpful discussions before and during this project. We thank John Philippi for support with electronic triggering. We thank Johannes Hohlbein, Mattia Fontana, Katarzyna Tych, Mahipal Ganji, and David Dulin for helpful comments on the manuscript. Parts of the analysis code used herein was co-developed in Thorsten Hugel’s lab by previous lab members (including SS). We are thankful for this contribution.

## Data & code availability

Raw data and new code created in this study is available for download at zenodo: 10.5281/zenodo.5972932.

