## Supplementary information for "Can DyeCycling break the photobleaching limit in single-molecule FRET?"

Vermeer, Schmid

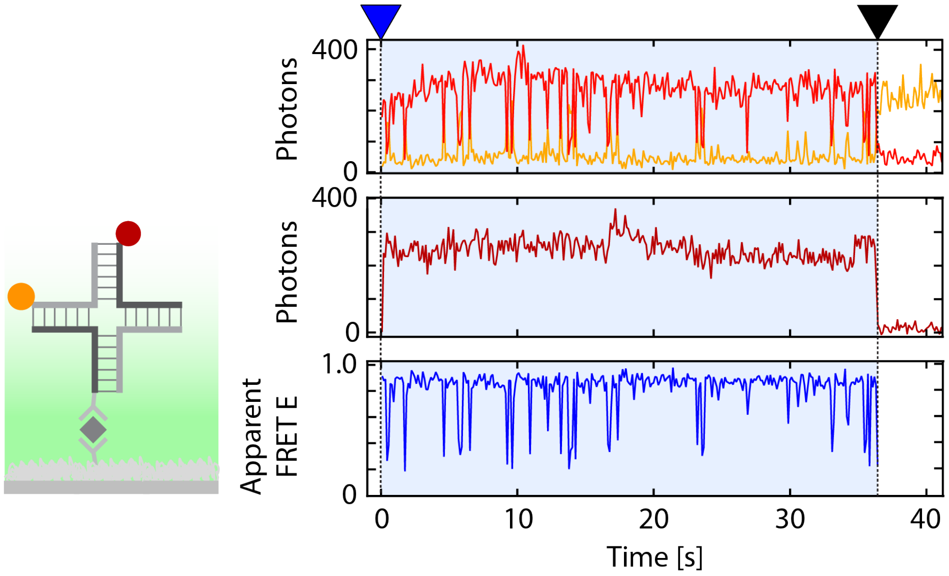


***Figure S1: Covalently labeled Holliday junction under identical conditions as the DyeCycling measurement.*** *Left: Illustration of a Holliday junction with a covalently attached donor (orange) and acceptor fluorophore (dark red). Right: Fluorescence and apparent FRET trajectories show the conformational changes of a Holliday junction as anticorrelated spikes in the top panel, and spikes in the bottom panel. Colors: donor: orange, FRET-sensitized acceptor: red, directly excited acceptor: dark red, apparent FRET efficiency (FRET E): blue. Blue triangle: start of the FRET regime (blue shading). Black triangle: acceptor photobleaching and end of the FRET regime.*

***Table S1. Holliday junction oligos and modifications.***

| **Identifier** | **Sequence** | **Modification** |
|---|---|---|
| HJ1 | 5’-CCC TAG CAA GCC GCT GCT ACG G-3' | |
| HJ2 | 5’-CCG TAG CAG CGA GAG CGG TGG G-3' | |
| HJ3 | 5’-CCC ACC GCT CTT CTC AAC TGG GGA TTA TCG CCG TTC T-3' | 5' biotin, C6-amino linker |
| HJ4 | 5’-CCC AGT TGA GAG CTT GCT AGG GCA TTC TCC TGT G-3’ | |
| DC1 | 5’-AGG AGA A**T**G-3' | dT-Atto647N, C6-amino linker |
| DC2 | 5’-GCG ATA A**T**C-3' | dT-Atto550, C6-amino linker |
